# Physical limits to acceleration of chemical reactions inside phase-separated compartments

**DOI:** 10.1101/2022.05.05.490822

**Authors:** Jeremy D. Schmit, Thomas C. T. Michaels

## Abstract

We present a theoretical analysis of phase separated compartments as a means to facilitate chemical reactions. We find that the attractive interactions that concentrate reactants within the dense phase inhibit reactions by lowering the chemical potential and mobility of the reactants. Therefore, condensed phases are only beneficial if mobility in the condensed phase can be maintained. This can be achieved in multi-step reactions, where the proximity between enzymatic steps results in higher efficiency with less unreacted substrate, but does not increase the reaction rate. Alternatively, mobility can be maintained if recruitment to the condensed phase is driven by multiple attractive moieties that can bind and release independently. However, the spacers necessary to ensure independence between stickers are prone to entangle with the dense phase scaffold. We find an optimal sticker affinity that balances the need for rapid binding/unbinding kinetics and minimal entanglement. Reaction rates can be accelerated by shrinking the size of the dense phase with a corresponding increase in the number of stickers to enhance recruitment.

---

In recent years many cellular structures, including nucleoli, Cajal bodies, stress granules and P-bodies, have been found that form by the spontaneous condensation of biomolecules [1–4]. These structures, termed membraneless organelles (MLOs) or liquid condensates, form via liquid-liquid phase separation and typically contain tens to hundreds of different molecular components that are enriched in concentration relative to the surrounding environment [1–4]. Usually, only a small subset of these components, called “scaffolds”, are necessary to drive condensation; the remaining components are classified as “clients”, which are not essential for condensation, but are recruited to the condensate by interactions with the scaffolds [4–6].

The biological function of MLOs remains a subject of intense research [3]. While experiments have revealed a variety of functions in well-studied systems [7–10], in most cases the function has not been established. Proposed functions include environmental sensing [8], stress response [9], signalling control [10], concentration buffering [11], and, in general, compartmentalization of biomolecular reactions in the cell [12]. In particular, it has been noted that MLOs could assist reaction kinetics by creating subcompartments with locally increased reactant concentrations [13–17]. However, the increased concentration also has the inhibitory effect of higher viscosity due to crowding effects [1, 18, 19] and reduced client mobility due to interactions with the scaffold. It therefore remains unclear to what extent MLOs can act as reaction crucibles to accelerate biochemical reactions.

Here we use a simple theoretical model to explore the effectiveness of MLOs at accelerating chemical reactions between clients. We find that the attractive interactions that recruit molecules to a MLO necessarily impair reaction dynamics by reducing diffusion. While MLOs are usually detrimental to reaction rates, we find two cases where they are beneficial. First, MLOs can achieve elevated concentrations without reducing diffusion when the substrate for a reaction is produced within a MLO, i.e., for multi-step reactions. Secondly, the strong concentration dependence of reaction orders ≥ 3 can overwhelm the inhibitory effects of client binding to the scaffold. These results shed light on the physical limits on how much phase-separated compartments can affect biochemical reactions.

## Model

To investigate the crucible hypothesis we consider a generic enzymatic reaction *nA* + *B* → *B* + *C* which describes the catalytic production of *C* from substrates, *A*, and a catalyst, *B*, that are clients of a condensate; *n* is the reaction order with respect to substrate concentration. The substrates and enzymes are present at concentrations *A*_*i*_ and *B*_*i*_ inside the condensate and *A*_*o*_ and *B*_*o*_ in the surrounding environment, respectively. The ratios *P*_*A*_ = *A*_*i*_*/A*_*o*_ and *P*_*B*_ = *B*_*i*_*/B*_*o*_ between inner and outer concentrations are the partitioning degrees of *A* and *B* molecules. The condensate, which is formed by phase-separation of a polymer (scaffold), has volume *V*_*i*_, while *V*_*o*_ denotes the volume of dilute phase, where *V*_tot_ = *V*_*i*_ + *V*_*o*_ is the total volume. To capture the interaction between the clients and the condensate scaffold, we further divide the interior concentration into soluble fractions, *A*_*is*_ and *B*_*is*_, and a fraction that is bound to the condensate scaffold, *A*_*ib*_ = *A*_*i*_ − *A*_*is*_ and *B*_*ib*_ = *B*_*i*_ − *B*_*is*_. For simplicity, we assume that the scaffold concentration in the dilute phase is negligible compared to that in the condensed phase, such that all *A* and *B* molecules in the outer phase are in the soluble state.

### Mobility loss upon binding impairs reaction rates

We first consider the situation where the species bound to the condensate scaffold are immobile. This means that reactions are limited to collisions with soluble species. This is known as the Eley-Rideal mechanism [20]. Examples of this mechanism include situations where the enzymes are covalently bound to the scaffolds [15, 16] or the clients are recruited by specific binding modules [21].

Under these circumstances, the reaction rate in the volume *V*_*o*_ surrounding the condensed phase is 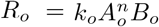, where *k*_*o*_ is the rate constant in the outer phase. Inside the condensate we impose the requirement that, at most, only one species can be bound for the reaction to proceed. This gives the reaction rate

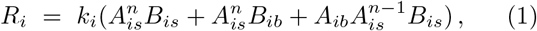

where *k*_*i*_ is the rate constant in the inner phase. The volume averaged rate is 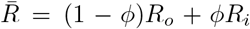, where *ϕ* = *V*_*i*_*/V*_tot_ is the volume fraction of the dense phase. If the reaction rate constants are equal in the two phases, *k*_*i*_ = *k*_*o*_ = *k*, and the system is in diffusive equilibrium across the condensate interface (*A*_*o*_ = *A*_*is*_ and *B*_*o*_ = *B*_*is*_), then the average reaction rate in the two-phase system is

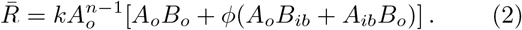

This should be compared to the well-mixed system with homogeneous concentrations 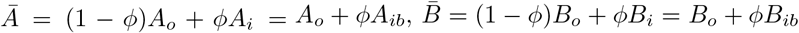 and total rate

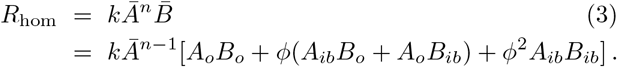

The rate *R*_hom_ of the homogeneous system is clearly greater than the total rate 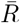 in the phase-separated system since *Ā > A*_*o*_ and 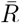 lacks the *ϕ*^2^ term. The missing term is due to the fact that molecule pairs that were immobilized in the phase-separated system are able to react in the homogeneous system. An important case is *n* = 1 and *A*_*ib*_ = 0, which describes a condensed enzyme and a substrate that does not bind the scaffold [15, 16]. In this case 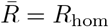. Therefore, the maximum achievable reaction rate is identical to that of a well-mixed system in the absence of condensate. Therefore, in the Eley-Rideal limit, condensate crucibles are either neutral or detrimental to reaction kinetics.

### Depletion of substrate within the condensate further inhibits reactions

The above analysis only considers the instantaneous rate at the beginning of the reaction. After the reaction begins there will be additional limitations due to diffusive transport between the condensate and surroundings and the exchange between bound and unbound states in the condensate. To understand these kinetic factors we consider the case where the enzyme is covalently linked to the scaffold so *B*_*is*_ = 0 and allow the *A* concentrations to vary with time. The kinetic equations for *n* = 1 are

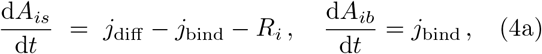

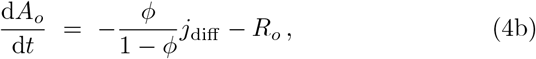

The concentration of soluble *A* particles within the condensate can change via three processes. The first term *j*_diff_ = *k*_diff_ (*A*_*o*_ − *A*_*is*_) describes the diffusive flux between the dense and dilute phases with *k*_diff_ = 3*D/r*^2^, where *D* is the diffusion constant of the *A* species and *r* is the condensate radius. The second term *j*_bind_ = *k*_on_[*A*_*is*_(*S*_*i*_ − *A*_*ib*_) − *K*_*D*_*A*_*ib*_] describes the binding and unbinding of *A* molecules to scaffolds in the condensate. We assume that the scaffold matrix provides binding sites at a concentration *S*_*i*_ of which *S*_*i*_ − *A*_*ib*_ are available to bind soluble *A* at a rate constant *k*_on_. The rate at which *A* are released from the scaffold, *k*_off_, is expressed in terms of the dissociation constant *K*_*D*_ = *k*_off_ */k*_on_. The third term in Eq. 4a describes *A* particles reacting with *B* at a rate *R*_*i*_. Eq. 4b describes the concentration of *A* particles outside of the condensate, which can change by either diffusive exchange with the condensate or reactions with *B* with rate *R*_*o*_. The diffusion term in Eq. 4b depends on *ϕ* reflecting the unequal volumes of the phases. Finally, the third equation describes the concentration of *A* particles that are bound to the condensate.

In the SI we compare the time required to consume half of the substrate for phase-separated vs. homogeneous systems. In all cases we find that the phase-separated system is slower than the homogeneous system and the reaction rates most closely approach the homogeneous limit when the rate constants are tuned to minimize binding to the condensate. This is true irrespective of whether the reaction is initiated with substrate homogeneously distributed throughout the system or when the reaction starts with substrate bound to the condensate.

### Condensates make multi-step reactions more efficient

We next consider two mechanisms by which condensates can increase the concentration of substrate while maintaining the mobility necessary for reactivity. The first mechanism is when substrate is produced within the condensate. For example, the production of ribosomes within the nucleolus requires multiple steps including RNA transcription, covalent RNA modification, RNA folding, and assembly with ribosomal proteins [22]. Sequential reactions can be handled by introducing a source term, *σ*, to Eq. 4. This addition results in two kinetic regimes. At short times, *t σ <* (*ϕA*_*i*_(0) + (1 − *ϕ*)*A*_*o*_(0)), the substrate generated by the source is negligible compared to the substrate present at *t* = 0 and the behavior is similar to Eq. 4 (Fig. 2). However, at long times the production of new substrate dominates and the system approaches a steady state solution in which the condensate plays a beneficial role. Since the localization is achieved by the production of substrate, we can understand the enhancement by neglecting the binding terms

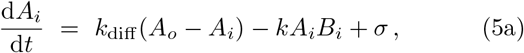

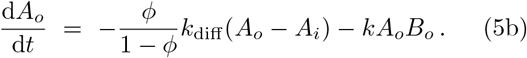

**FIG. 1:**
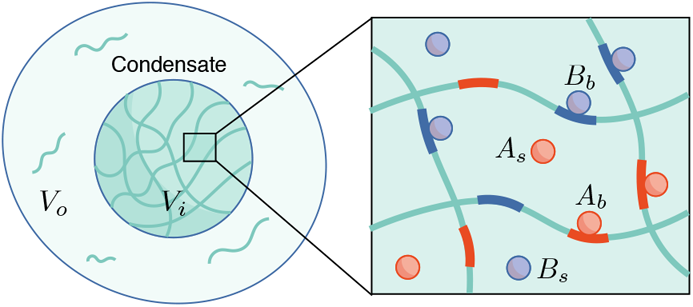
Schematic representation of model. A system of total volume *V*_tot_ contains a condensate of volume *V*_*i*_, which is formed by phase-separation of a polymer (scaffold) with respect to the surrounding cyto- or nucleoplasm. The system contains molecules *A* and *B* which undergo an enzymatic reaction and are clients of the condensate. The scaffold-client interaction is described in terms of sites on the polymer that *A* and *B* molecules can bind to. The concentrations of *A* and *B* molecules is therefore divided into soluble fractions, *A*_*s*_ and *B*_*s*_, and fraction bound to the condensate scaffold, *A*_*b*_ and *B*_*b*_.

**FIG. 2:**
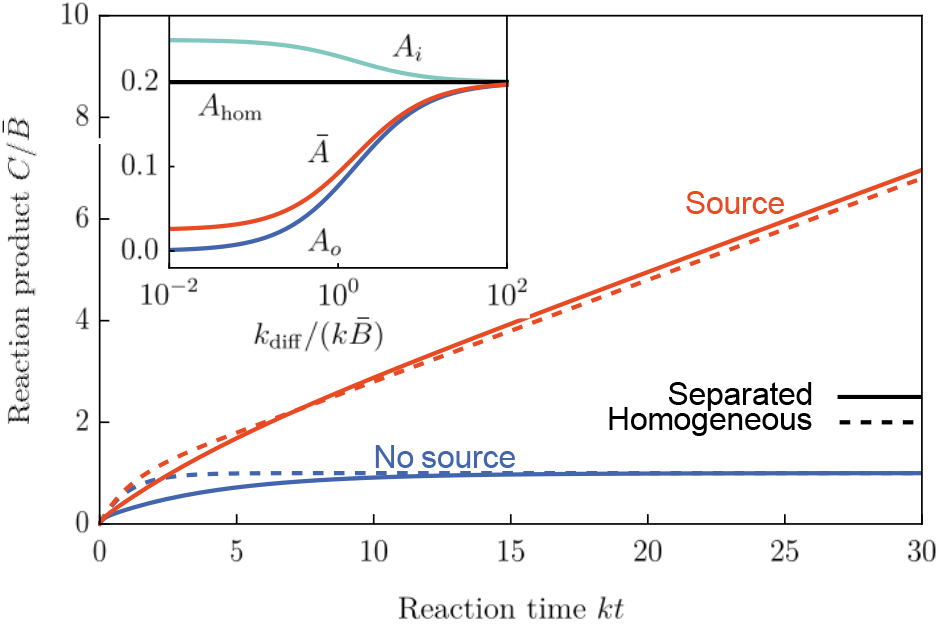
When the supply of substrate is fixed (blue lines) recruitment of enzymes to a condensate reduces the rate of product formation because the diffusive transport of substrate is limiting. When substrate is produced within the condensate (red lines) the system initially behaves like the fixed substrate system where the condensate is detrimental before converging to a steady state where the condensate is somewhat beneficial. Inset: The steady state concentrations of substrate in the dilute phase (*A*_*i*_), dense phase (*A*_*o*_), and total system (*Ā*) all converge to that of the homogeneous system (*A*_hom_) when the diffusion rate is very fast.

The steady-state solution to these equations is

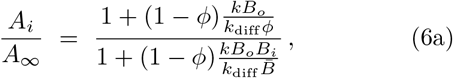

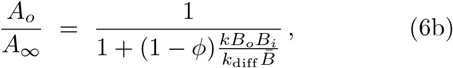

where 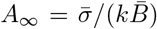 is the homogeneous solution limit (equivalent to *k*_diff_ → ∞) and 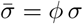. From this we can see that the outside concentration *A*_*i*_ is always less than the homogeneous case while inside concentration *A*_*o*_ is always greater. To see the benefit of the condensate, we look at the average concentration of *A*

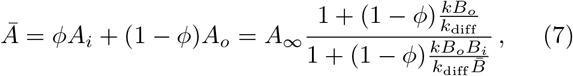

which shows that *Ā* is always less than the homogeneous limit *A*_∞_, since 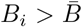 (Fig. 2 inset). Importantly, the condensate does not affect the overall reaction rate, which is fixed by the steady-state condition. However, the condensate ensures that there is less unused substrate. This is particularly useful for limiting the quantity of intermediate states in high-volume reactions like ribosome synthesis as well as situations where intermediate states are prone to pathogenic aggregation.

### Small condensates optimize reactant concentrations

Thus far we have assumed that molecules are immobile when bound to the condensate network. However, condensate scaffolds are not uniformly attractive and, instead, have discrete sticky moieties responsible for condensation and client recruitment [23, 24]. The clients can remain mobile while bound if they have the same sticker-and-spacer architecture as the scaffolds because independent stickers allow some parts of the molecule to move while other parts remain bound. In the following we allow for mobility in the dense phase and show that condensates can be beneficial for promoting reactions provided the reaction order is sufficiently large.

When the interior and exterior rate constants, *k*_*i*_ and *k*_*o*_ differ the volume averaged reaction rate is

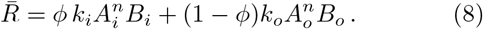

We assume that the enzyme and substrate are recruited to the condensate by equivalent interactions resulting in identical partition coefficients *P* = *A*_*i*_*/A*_*o*_ = *B*_*i*_*/B*_*o*_. Using the conservation of mass condition Eq. 8 can be written

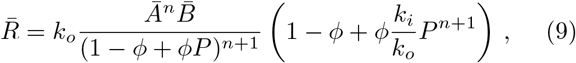

which we can optimize with respect to *ϕ* to yield

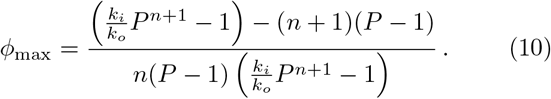

The optimal volume fraction is inversely proportional to the reaction order *n*. This is because large reaction orders are so sensitive to concentration that it is more beneficial to have a small reaction volume that is highly saturated than to have a larger reaction volume at lower concentration. A small condensate volume ensures that the dilute phase is minimally depleted, which for fixed *P* maximizes the concentrations in the dense phase.

### The optimal sticker affinity is a tradeoff between recruitment and mobility

While the sticker/spacer structure allows the client to move without detaching from the condensate, it comes at a price because the spacers that allow the stickers to bind/unbind independently are prone to entangle with the scaffold. As a result, the client molecules will transition from Stokes diffusion outside the condensate to reptation diffusion inside. Thus, if a certain level of recruitment is required, there is a tradeoff between a small number of stickers that bind tightly or a large number of weak stickers that are highly entangled. The optimum between these extremes can be determined as follows.

The first step is to determine how strongly a client is recruited to the condensate. The free energy change for a molecule with *M* stickers entering the condensate is (see SI)

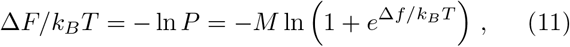

where Δ*f* is the difference in free energy between the bound and unbound states for a sticker and spacer.

The diffusion constant for a polymer undergoing reptation dynamics will be proportional to 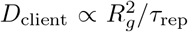 indicating that the polymer has moved a distance proportional to its radius of gyration *R*_*g*_ ∝ *L*^1*/*2^ during a reptation time, *τ*_rep_. The reptation time is the time required for the polymer to exit a tube of length *L* diffusing in the direction parallel to its backbone *τ*_rep_ ∝ *L*^2^*/D*_*‖*_, where *D*_*‖*_ describes diffusion in the direction parallel to the elongated direction. The parallel diffusion will be inversely proportional to the length of the polymer *D*_*‖*_ ∝ 1*/L*, where the proportionality constant is the diffusion constant of a chain subunit. In this context the repeating unit has contributions from spacer and the adjoining sticker. We assume that, for the sticker-spacer unit to move through the network, the sticker must be in the unbound state. Therefore, we have

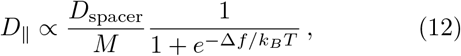

where we have used *L* ∝ *M*. Thus, we find

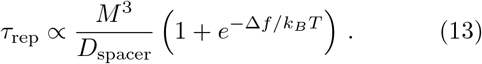

After *τ*_rep_ the polymer will exit a tube of entanglements, which requires the molecule to move by a distance 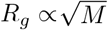. Therefore the diffusion constant is

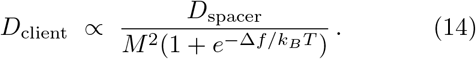

Using Eq. 11 we optimize *D*_client_ with respect to *M* holding the total affinity, Δ*F*, fixed. This yields *A/M* = 2 or 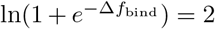. The optimal binding free energy for a sticker, including the sticker-sticker interaction and the conformational entropy loss of the spacer, is therefore about 2*k*_*B*_*T*. This value allows the sticker to bind and release frequently enough for the molecule to diffuse, but provides enough affinity the molecule does not become too long and entangled.

### Reaction rates are enhanced for high order reactions

With this result we can determine the optimal level of client recruitment to maximize the reaction rate. We assume that *k*_*i*_ is proportional to the client diffusion constant so *k*_*i*_*/k*_*o*_ = *α/M*^2^, where *α* is a constant accounting for the transition between Stokes and reptation dynamics. Inserting this into Eq. 9 and using the fact that the optimal sticker affinity gives 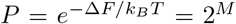 (Eq. 11), we find that the reaction rate is greatest when the number of stickers is equal to the largest root of (see SI)

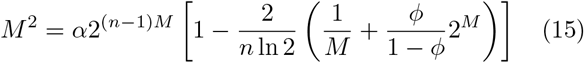

which can be approximated by *M* ≃ ln[*n*(1 − *ϕ*) ln 2*/*2*ϕ*]*/* ln 2. If *n* is small or *ϕ* is large there are no solutions to Eq. 15 indicating that recruitment to the condensate is detrimental to the reaction rate. However, large *n* and small *ϕ* favor highly partitioned systems with *M >* 10.

Fig. 3 shows 𝒪(10^3^) enhancement for *n* = 3, which shows why condensates are prone to the nucleation of aggregates [25] and why they are useful for the assembly of large complexes like viral capsids [26, 27]. Conversely, for small *n* it is difficult to get any rate enhancement. Fig. 7 shows that second order reactions require *ϕ <* 10^−3^ and recruitment affinities of |Δ*F*| *>* 10*k*_*B*_*T* to achieve even modest enhancements (less than 10-fold). This regime may not be physically realistic because the high concentration of clients will compete for available stickers, voiding the independent molecule approximation of Eq. 11, and limiting the achievable values of *P* (unless the reactants have very low copy numbers).

**FIG. 3:**
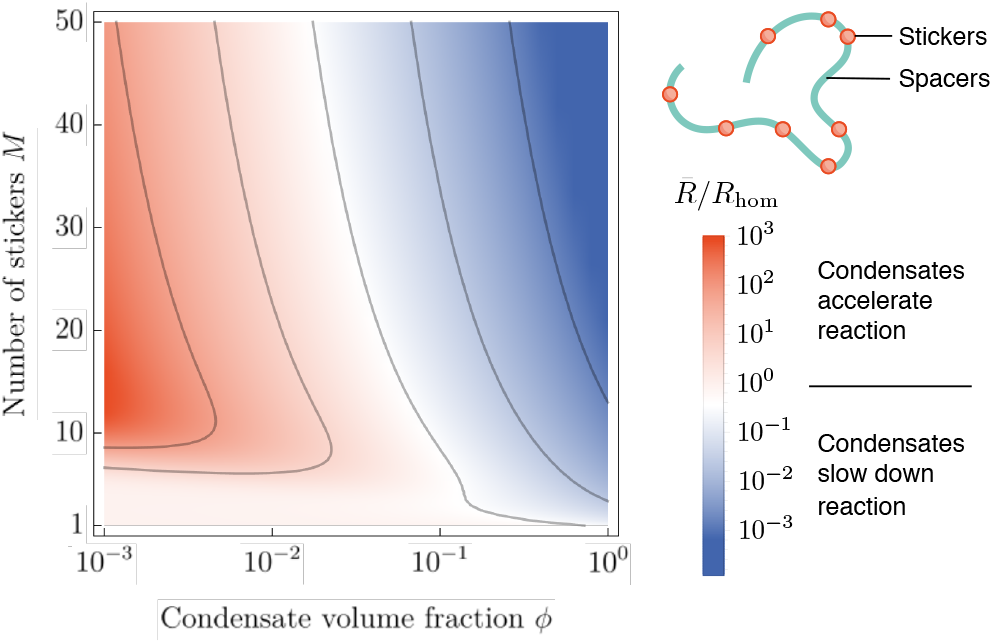
Ratio of the reaction rate (Eq. 8) for a separated system to a homogeneous system (Log_10_ scale) as a function of the condensate volume fraction *ϕ* and the number of 2 *k*_*B*_*T* stickers. When *ϕ* is sufficiently small there is an optimal number of stickers on the order of *M* ≃ 10 that maximizes the reaction rate. *n* = 3, *α* = 0.3

And important case that is not explicitly covered by our calculations is that of electrostatic coacervates. We expect that these will behave similarly to our reptation calculation, although the smooth charge distribution on molecules like RNA violates our approximation of independent stickers. Another complication is that the neutralization afforded by complementary charged scaffolds will perturb the conformational ensembles and stabilize compact and/or double stranded states [13, 14, 17].

In conclusion, we have shown that the interactions necessary to recruit reactants to phase separated compartments necessarily impairs mobility, and hence reactivity. This mobility loss may, nevertheless, be acceptable in situations with multiple sequential reactions or high order reactions.

## Acknowledgments

We acknowledge support from Institute for the Physics of Living Systems, University College London, and National Institutes of Health grant R01GM141235.

## SUPPLEMENTARY INFORMATION

### NUMERICAL SOLUTION OF RATE EQUATIONS WHEN BOUND REACTANTS ARE IMMOBILE

Fig. 4 plots the condensate enhancement factor, which we define as the time required for the reaction to consume half of the *A* particles in the homogeneous system divided by the separated system. The former quantity is computed from 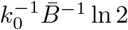, while the latter quantity is determined by numerical solution of Eqs. 4a. In all cases the enhancement factor is less than one, indicating that the presence of the condensate has an *inhibitory* effect on the reaction rate. This is true whether the system is initialized from a homogeneous state, *A*_*o*_ = *A*_*is*_ = *Ā* and *A*_*ib*_ = 0 (Fig. 4), or if the system is started from a state where the *A* particles have reached an equilibrium binding to the condensate before reaction is started (data not shown).

**FIG. 4:**
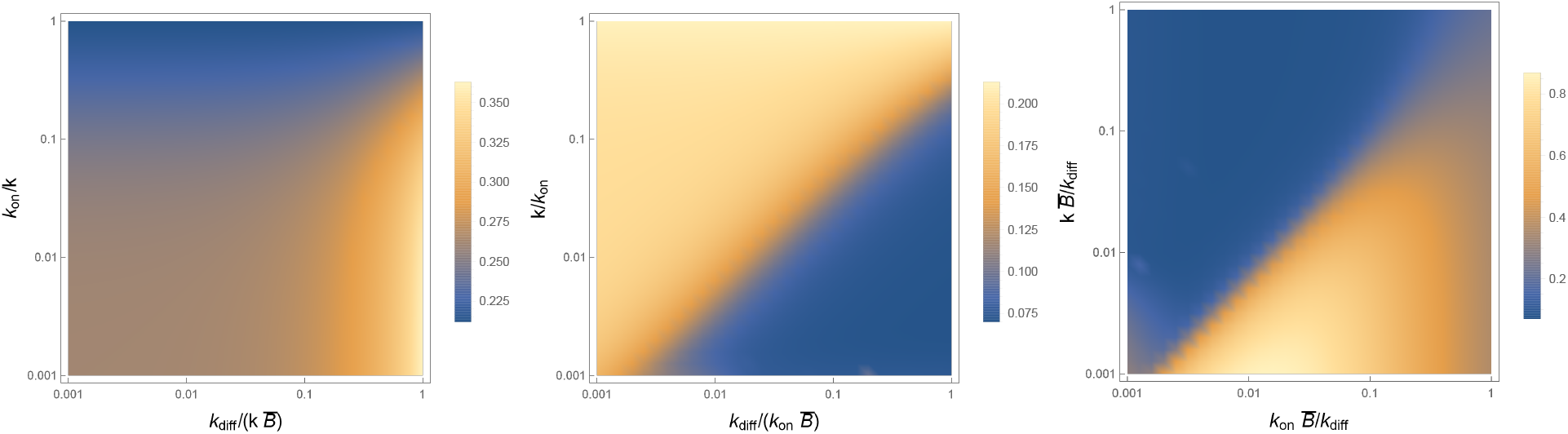
Plot of the condensate enhancement factor as a function of the rate constants in Eqs. 4a. The enhancement factor compares the time to consume half of the reactant in the homogenous system to the separated system. Ratios less than one, which is the case in all conditions in the figure, indicate that it takes longer for the reaction to proceed in the separated system. In each panel the fastest rate constant is held fixed (*k* in the left panel, *k*_on_ in the middle panel, and *k*_diff_ in the right panel) and the other two are varied over three orders of magnitude. See text for a brief description of each panel. 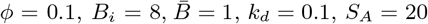

#### Fast reactions

When the reaction rate is faster than the binding or diffusion timescales the condensed system most closely approaches the rate of the homogeneous system when *k*_diff_ is large and *k*_on_ is small. Large *k*_diff_ minimizes the inhibitory effect of transporting substrate to the enzymes in the condensate, while small *k*_on_ allows the substrate to react with immobile enzymes before it binds to the condensate.

#### Fast binding

When binding to the network is very fast, the way to keep substrate in the unbound, reactive state is to keep the soluble substrate concentration low (below *K*_*D*_). This occurs when *k ≫ k*_diff_ so that the reaction consumes substrate as soon as it enters the condensate.

#### Fast diffusion

When the diffusion rate is much faster than the reaction or binding rates, the inhibitory effect of substrate transport is negligible and the system approaches the reaction rate of the homogeneous system, provided the reaction consumes the incoming substrate faster than it can bind. If the reaction rate is too fast, however, diffusion can become limiting again. The optimal reaction rate in the right panel of Fig. 4, where the reaction proceeds at ∼ 80% of the rate of the homogeneous system, occurs when *k* is large enough to out-compete binding, but not so large that diffusion becomes limiting.

### RECRUITMENT EQUILIBRIUM

In this section we will calculate the concentration of polymer-like clients within a scaffold network. The extent of recruitment is determined by the condition that the chemical potential of the clients must be the same in the dilute and concentrated phases. The partition function for *N* clients is given by 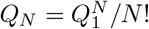. This expression is valid in the dilute phase, but not necessarily in the dense phase. If recruitment to the dense phase is strong enough, it is necessary to account for interactions between the clients. These interactions will result in steric collisions and/or competition for scaffold binding sites. Both of these interactions are repulsive and will lead to a lower level of recruitment than what we compute here. Therefore, this calculation represents an upper limit on the ability of a condensate to facilitate reactions.

The chemical potential of the clients is

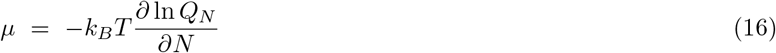

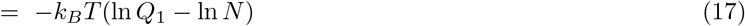

The next step is to evaluate *Q*_1_ which we write as a product of position and conformation states *Q*_1_ = *Q*_pos_*Q*_conf_. To evaluate the position partition function we choose one end of the polymer as a reference point and sum over all possible positions for this point of the polymer. Neglecting boundary effects, this integration yields the volume of the relevant phase divided by a microscopic parameter *V*_1_ that sets the volume per state (similar to the Debye wavelength).

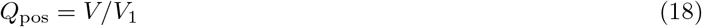

Inserting this into the chemical potential we have

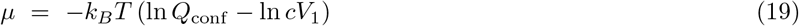

where *c* = *N/V* is the client concentration.

To evaluate *Q*_conf_ we assume that the molecule consists of *M* stickers and *M* + 1 spacers (i.e., both ends of the molecule teminate with a spacer). We further assume that each spacer is long enough, and that the number of scaffold binding sites is large enough, that each spacer is statistically independent. Each spacer has a fixed starting point that is determined by either the state in the position integration (for the first spacer) or the end point of the previous spacer (for the next *M* spacers). The partition function for each spacer consists of a sum over all possible end points given the fixed starting point. In the dilute phase, where we can neglect binding, each spacer is identical so

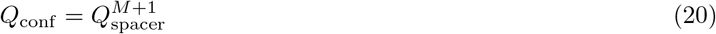

where *Q*_spacer_ accounts for the Gaussian distribution of polymer states for fixed starting and ending points

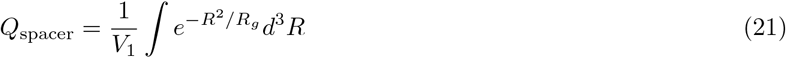

where *R* and *R*_*g*_ are the end-to-end vector and the radius of gyration of the spacer, respectively.

In the dense phase we need to account for the fact that the first *M* spacers will have their conformational statistics perturbed by binding to the scaffold. However, the (*M* + 1)th spacer, which does not contain a sticker is unperturbed by the scaffold (we neglect crowding effects). Therefore, in the dense phase we have

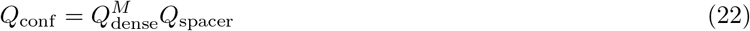

The sum over states for the *M* spacers with stickers is similar to the dilute phase in that the starting point is fixed by either the position integration or the previous spacer. We account for the effect of the scaffold by splitting the integration into terms that result in either a free end or sticker-scaffold binding

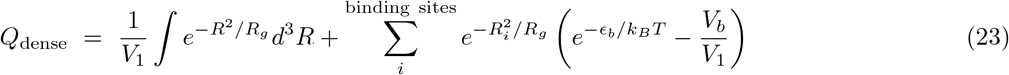

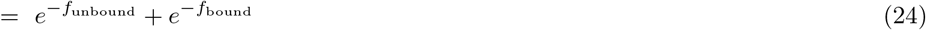

In the second term *ϵ*_*b*_ is the affinity of the sticker-scaffold binding interaction and *R*_*i*_ is the distance between the *i*-th scaffold binding site and the beginning of the spacer. In writing the unbound integral we have over-counted the volume accessible to the end of the spacer because each binding site on the scaffold will be surrounded by a volume *V*_*b*_ that results in a bound state. This over-counting is corrected by the *V*_*b*_*/V*_1_ term of the sum over binding states, which assumes that the *V*_*b*_ is small enough that *R*_*i*_ does not change appreciably within this volume (Fig. 5).

**FIG. 5:**
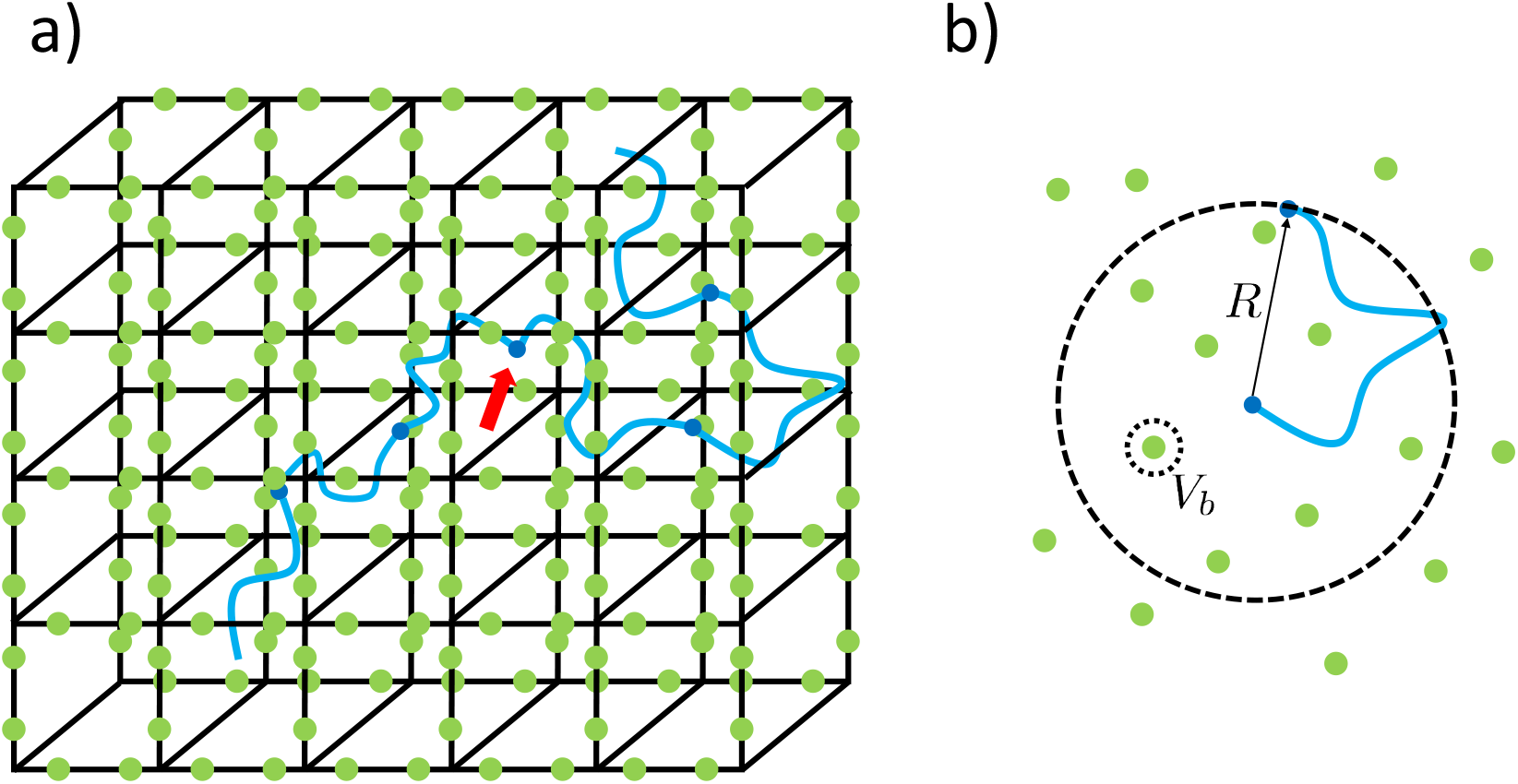
Schematic of the geometry in the calculation of the client free energy. (a) The calculation describes a polymer-like client (blue) entangled with a scaffold matrix (black lines). Stickers on the client (dark blue dots) can interact with attractive patches (green dots) on the scaffold. The spacers are long enough, and the binding patches abundant enough, that the binding states of adjacent stickers are approximately adjacent. The figure shows a client with *M* = 5 stickers. Four of these stickers are bound to the scaffold but the middle one is unbound (red arrow). This independence allows the client to gradually move through the network. (b) The partition function for each spacer involves an integration over the end to end vector *R*. This integration will include states where the sticker at the end is unbound (as shown) and states with it bound to the scaffold binding sites (green dots). Each binding site is surrounded by a volume *V*_*b*_ within which the sticker is considered bound.

At equilibrium we have *μ*_dilute_ = *μ*_dense_ which gives

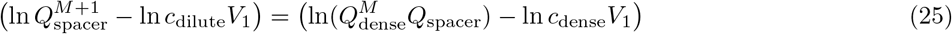

This expression can be rearranged as follows

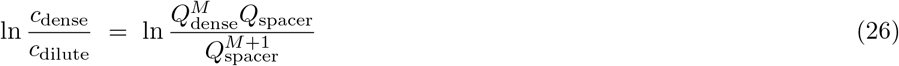

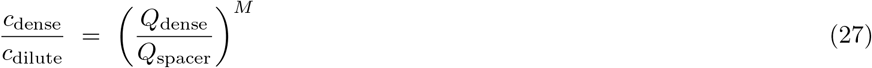

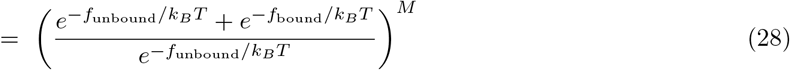

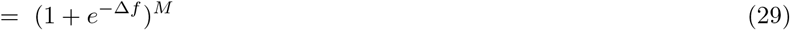

where Δ*f* = *f*_bound_ − *f*_unbound_.

Our next task is to connect these free energies to the enrichment/depletion due to condensate formation. As defined in the text, the partition coefficient is given by

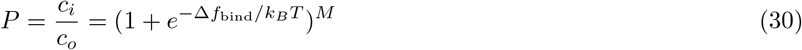

where *c*_*i*_ and *c*_*o*_ are the concentrations in the dense and dilute phases, respectively.

### OPTIMAL BINDING AFFINITY

Increasing the affinity of a client to the condensate comes at the cost of mobility. Here we determine the optimal affinity that balances the increase in concentration with the loss in mobility. We start with the volume averaged reaction rate (Eq. 8)

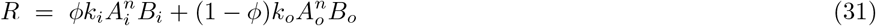

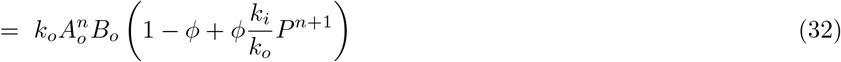

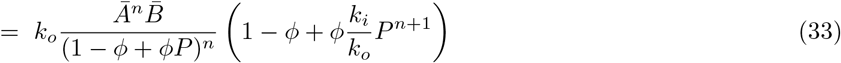

where the second line assumes that both *A* and *B* molecules have optimal recruitment characteristics so Δ*F*_*A*_ = Δ*F*_*B*_ and hence *P* = *P*_*A*_ = *P*_*B*_. The third line follows from the conservation of mass condition 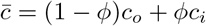.

Increasing the affinity of clients to the network has three effects on the reaction rate. First, it reduces the dilute phase concentration, which reduces the reaction rate. Second, it increases the dense phase concentration, which increases the rate. Third, it decreases the mobility of reactants in the dense phase, which reduces the reaction rate. The first two effects are captured by the *P* dependencies in Eq. 33, but the calculation in the main text shows that the optimal sticker affinity is given by ln 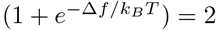, so *P* = 2^*M*^ in the best case scenario.

The mobility effect is captured in the *k*_*i*_*/k*_*o*_ factor. We assume that the reaction rate inside the condensate *k*_*i*_ is proportional to the diffusion constant of the clients. The diffusion constant is proportional to 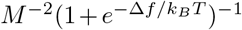 (Eq. 14), but the term in parentheses is just another factor of 2 that can be absorbed into the constant of proportionality.

Therefore, we write

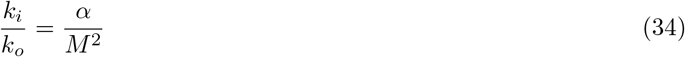

where *α* is a constant of proportionality (that will have minimal effect on the results).

Now we insert Eq. 34 and partition coefficients into Eq. 33

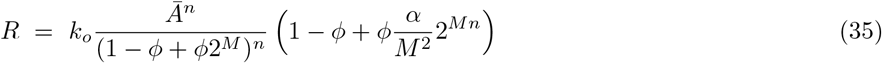

This expression has a (non-physical) divergence at *M* = 0 and monotonically decreases at large *M*. There are two possible behaviors in-between, it can monotonically decrease, or there can be a local minima followed by a local maximum (Fig. 6).

**FIG. 6:**
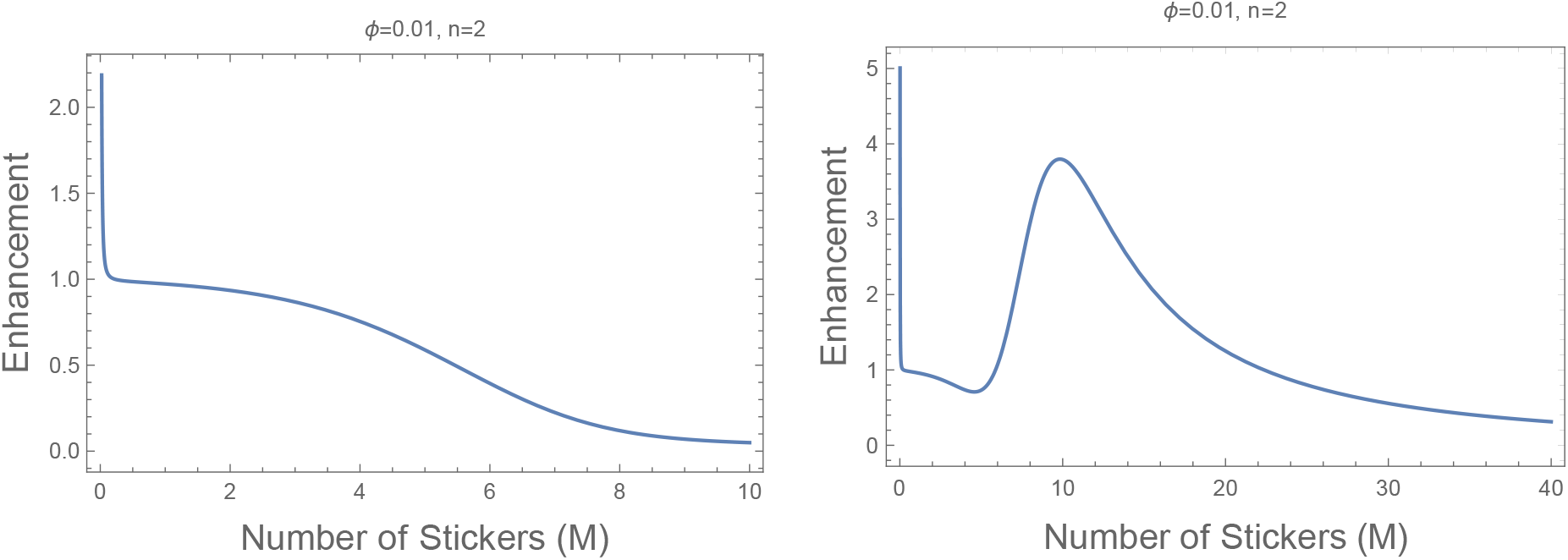
The reaction rate can be either a monotonically decreasing function of the number of stickers *M*, or it can have a pair of local extremes.

**FIG. 7:**
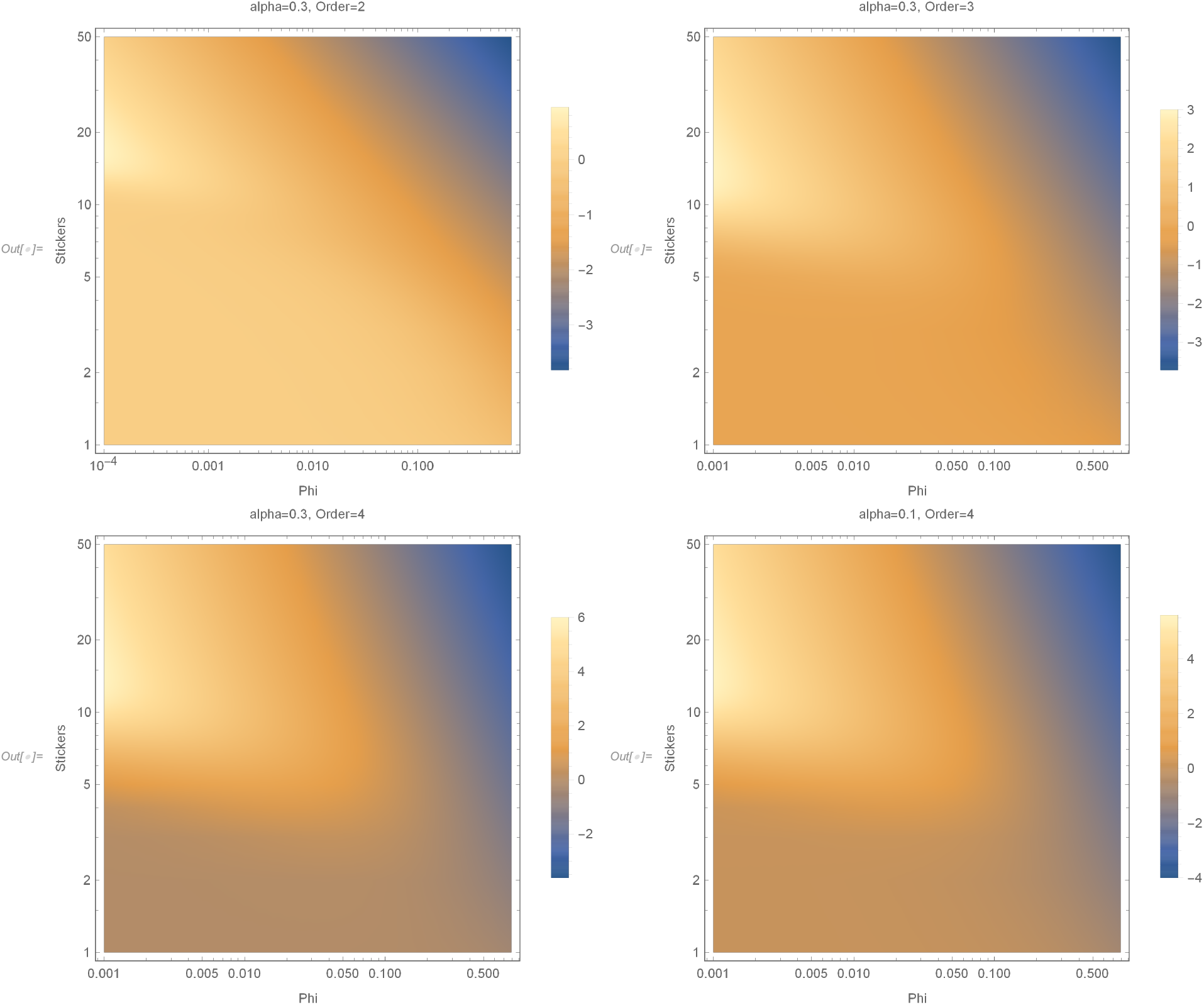
Log_10_ of the reaction rate as a function of the volume fraction and the number of stickers. The reaction order makes a big difference but, *α*, the constant of proportionality between Stokes and reptation diffusion, does not.

To determine if there is an enhancement due to recruitment, we take the derivative of the rate with respect to *M*

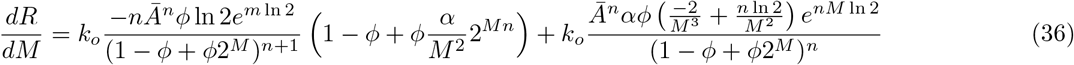

To find the roots of the equation we find a common denominator and set the numerator equal to zero

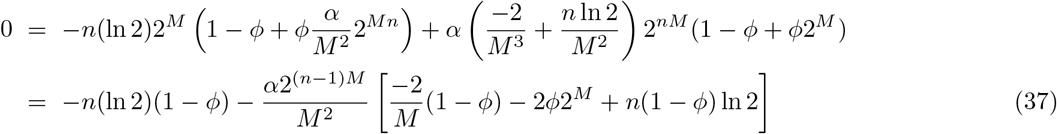

To analyze this equation, we rearrange it as follows

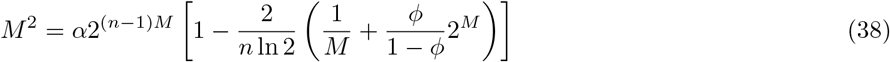

This is a transcendental equation that can be solved graphically. We are looking for intersections between the left side and the right side. The physically relevant region is *M >* 1 but mathematical intuition can be gained by starting at *M* = 0.

- The left side starts at 0 at *M* = 0 and diverges to +∞ as *M* → +∞.
- The right side tends to −∞ at *M* = 0 (dominated by the 1*/M* term) and also tends to −∞ as *M* → +∞ (dominated by the 2^*M*^ term).
- The right side can only intersect the left side if the term in square brackets is positive. Despite the fact that the left side increases rapidly with *M*, the right side diverges even faster due to the 2^(*n*−1)*M*^ prefactor. Therefore, even a slightly positive value in the square bracket is likely to give a root.
- The biggest unknown in this equation is the constant *α*. While this parameter has a strong effect on the maximum rate that is achievable within the dense phase, it has a very weak effect on the presence or location of the maximum. That is because the right side of the equation will be nearly vertical at the two roots, meaning that even an order of magnitude change in *α* will only change the location of the roots by *M* ± 1.
- The interesting root, representing the rate maximum, is close to the largest value of *M* for which the term in the round parentheses is less than 1 (including the prefactor). For larger values of *M* the 1*/M* term is negligible, so we are primarily interested in the second term. Therefore a good approximation for the optimal number of stickers, *M*, is given by

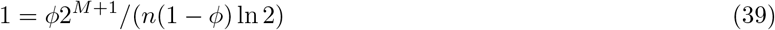

which can be rearranged to give

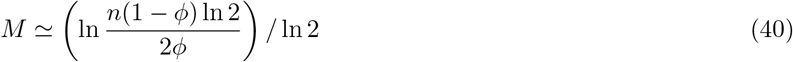

In searching for roots of Eq. 39, note that the 2^*M*^ factor grows very rapidly, but the factor of *ϕ* can be very small. For *ϕ* on the order of 10^−1^ to 10^−2^ the solutions will be in the range of *M* = 3 to *M* = 7. If the reaction order *n* is large, the peak value of *M* may be even larger than this.

## Notes

### Competing Interest Statement

The authors have declared no competing interest.

